# Efficient coding in working memory is adapted to the structure of the environment

**DOI:** 10.1101/2025.08.20.671058

**Authors:** Qiaoli Huang, Christian F. Doeller

## Abstract

Working memory (WM) relies on efficient coding strategies to overcome its limited capacity, yet how the brain adaptively organizes WM representations to maximize coding efficiency based on environmental structure remains largely unknown. In our study, participants remembered a sequence of gratings defined in a two-dimensional feature space where we manipulated directional consistency, revealing enhanced performance for structured (consistent direction) vs non-strutured (non-consistent direction) contexts, particularly for individuals with lower WM capacity. Magnetoencephalography analyses uncovered dissociable neural bases: consistent sequences engaged anterior temporal and medial frontal cortices for abstract directional representations during maintenance, while inconsistent sequences preferentially reactivated item-specific representations in parietal regions. These neural patterns predicted behavioral performance, establishing a neural efficiency principle wherein the brain adaptively switches between relational and item-based coding strategies, mitigating WM constraints. These findings advance our understanding of how structures shape WM organization, offering insights into cognitive flexibility and neural resource allocation in complex environments.

## Introduction

Working memory (WM), a fundamental cognitive function for temporary information maintenance, has a limited capacity that constrains humans to retain only a small amount of information at a time (Baddeley, 1992; Cowan, 2001; Miller, 1956). This limitation prevents accurate retention of all sensory inputs, necessitating efficient information compression to optimize WM storage and processing (Bates & Jacobs, 2020; Mathy & Feldman, 2012). Decades of research have demonstrated that leveraging statistical regularities among multiple stimuli enables efficient data compression and leads to superior WM performance (Brady et al., 2009, 2011; Kaiser et al., 2015; Lew & Vul, 2015; Mathy & Feldman, 2012). Aligning with these behavioral observations, efficient coding theory proposes that neural resource constraints drive the development of efficient (compact) representational codes optimized for stimulus statistics of the environment (Attneave, 1954; Bays et al., 2024; Ganguli & Simoncelli, 2014; Wei & Stocker, 2015). However, direct neural evidence remains scarce regarding how the brain optimally represents abstract structural information in WM to achieve such efficient coding.

An entirely separate research line in neuroscience on cognitive maps suggests that the brain creates spatial maps or map-like abstract representations by encoding relational information in a compact, readily accessible manner (Behrens et al., 2018; Bellmund et al., 2018; Epstein et al., 2017; O’Keefe, 1978; Tolman, 1948; Whittington et al., 2020). These maps exemplify efficient coding principles by preserving relational structure while discarding unnecessary details, thereby optimizing rapid retrieval and application of behaviorally relevant information. The hippocampal-entorhinal system, with its spatially tuned neurons, forms the neural basis of these cognitive maps (Buzsáki & Moser, 2013; Hafting et al., 2005; O’Keefe & Dostrovsky, 1971).

Recent advances reveal this system’s broader role in coding abstract relationships across diverse domains, including associative learning, social hierarchy inference, and prospective decision-making (Bao et al., 2019; Barnaveli et al., 2025; Constantinescu et al., 2016; Nitsch et al., 2024; Park et al., 2021; Theves et al., 2019; Theves et al., 2020; Viganò et al., 2023). Collectively, these findings indicate that cognitive maps might provide a more general mechanism for organizing relational information across multiple behavioral dimensions (Whittington et al., 2020). Building on this framework, we hypothesized that the brain employs similar neural coding principles to achieve efficient compression of multiple WM representations by extracting and utilizing their underlying abstract relationships.

A well-known feature of WM is its distributed representation across multiple cortical areas (Christophel et al., 2017), ranging from primary sensory cortices (Christophel et al., 2012; Harrison & Tong, 2009; Serences et al., 2009; Van Kerkoerle et al., 2017) to the frontal regions that code more abstract stimulus features, e.g., categorical information (Freedman et al., 2001; Vergara et al., 2016; Wallis et al., 2001). Recent studies, particularly evidence from intracranial recordings, demonstrate that the medial temporal lobe is also essential for WM, particularly under challenging conditions involving distractors, high memory load, or specific content types such as spatial-context information and abstract concepts (Daume, Kamiński, Salimpour, et al., 2024; Daume, Kamiński, Schjetnan, et al., 2024; Jeneson & Squire, 2012; Kamiński et al., 2017; Libby et al., 2014). These findings reveal a highly flexible system capable of representing diverse WM contents—from sensory details to abstract structure—across distinct brain regions. A key unresolved question, however, is how the brain dynamically prioritizes and optimizes specific representations to efficiently store multiple information across different contexts.

To address this, we experimentally manipulated the statistical regularities among WM stimuli, hypothesizing that systematic variations in these regularities would drive adaptive neural representations to optimize coding efficiency. We recorded whole-brain magnetoencephalography (MEG) signals while participants memorized a sequence of three gratings varying along two continuous dimensions: orientation and frequency (Figure 1A). Each grating can thus be described as a point within a two-dimensional (2D) feature space. Critically, we manipulated the alignment of vector directions between consecutive items in this 2D feature space. In the consistent direction (CD) condition, all three items formed a collinear arrangement with identical inter-item vector directions, whereas in the inconsistent direction (ICD) condition, the vector direction shifted between the 1^st^-to-2^nd^ and 2^nd^-to-3^rd^ item pairs (Figure 1BC). Importantly, the first two items were held constant across conditions, enabling a direct comparison of how the brain optimally organizes identical stimuli under different statistical contexts. We predicted that the preserved directional regularity in the CD condition would allow efficient compression of multiple information through the extraction of shared relational codes, rather than requiring independent item storage, thereby circumventing WM capacity limitations.

**Figure 1.**
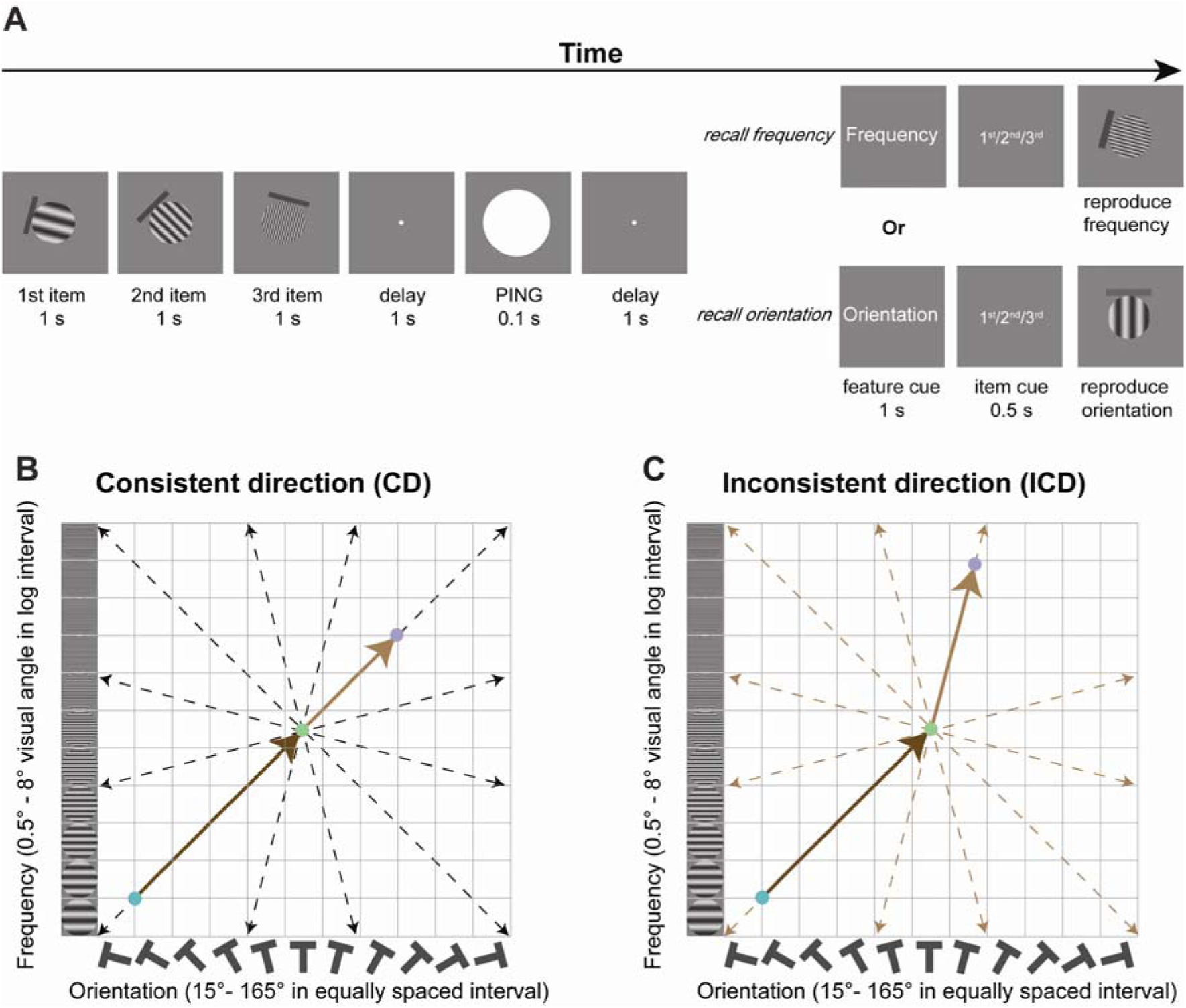
Experimental paradigm. **(A)** At the beginning of each trial, three gratings with different orientations and frequencies were sequentially presented for 1 s with 0.75 s interval, and participants were asked to memorize both the frequency and orientation of the sequence and later reproduce the cued domain (orientation or frequency) of the cued item (1^st^ or 2^nd^ or 3^rd^) by using mouse to continuously increase or decrease the corresponding value of the test stimulus. During the maintaining period, a high luminance white disk (PING stimulus) was briefly presented (0.1 s), aiming to perturb the memory network to enable the readout of memory-related reactivation. The three gratings were sampled from a continuous 2D feature space defined by frequency (ranging from 0.5° to 8° visual angle in log interval) and orientation (ranging from 15° to 165° in equally spaced interval). Unknown to participants, each item can be characterized as a single point in the 2D feature space. Based on the way to sample the three items, there were two conditions, consistent direction (CD) and inconsistent direction conditions (ICD). (B) The CD condition. The three sequential items were located on a line, with a random distance for consecutive items and a random starting point. There were 12 possible directions (15°, 45°, 75°, 105°, 135°, 165°, 195°, 225°, 255°, 285°, 315°, 345°). (C) The ICD condition. Across trials, the same 1^st^ and 2^nd^ items were applied in the ICD condition. The 3^rd^ item was sampled with the constraint that it was not in the same direction as the 1^st^ and 2^nd^ items. See one example of how the three sequential stimuli were randomly sampled within the continuous 2D feature space for the individual participants in supplementary figure 1.

## Results

### Experimental procedure and behavior performance

Thirty-three participants completed two MEG recording sessions. In each session, three gratings with different orientations and spatial frequencies sequentially appeared at the center of the screen (Figure 1A). Participants were asked to remember both features (i.e., orientation and spatial frequency) of all three gratings. Following a maintenance period, participants were asked to reproduce the randomly cued feature of the cued item. During the maintenance period, a high-luminance PING stimulus was briefly presented, aiming to perturb the “activity-silent” WM network (Panichello et al., 2024; Stokes, 2015) to allow access to the actively stored information (Huang et al., 2021; Wolff et al., 2017). We manipulated the sampling of gratings within the 2D feature space (orientation X spatial frequency). In the consistent direction condition (CD), the three items were collinear, where the vector direction from the 1^st^ to the 2^nd^ item matched that from the 2^nd^ to the 3^rd^ item in the 2D feature space. Conversely, in the inconsistent direction condition (ICD), the direction from the 1^st^ to the 2^nd^ item differed from that of the 2^nd^ to the 3^rd^ item. Critically, we implemented a paired trial design: CD and ICD trials shared identical 1^st^ and 2^nd^ items, with condition differences emerging only at the 3^rd^ item position. Therefore, the between-condition memory difference for the first two items reveals how effectively the brain leverages consistent directional structure to compress multi-item representations, as opposed to maintaining items independently under capacity-limited constraints.

Memory precision was quantified as calculating the standard deviation (SD) of the memory error (the difference between reported orientation (frequency) and correct orientation (frequency)) separately for each item. A higher SD value indicated lower memory precision. We aggregated performance across both feature domains and the first two items (see individual domain/item memory performance in supplementary figure 2). While the mean memory error reflects the systematic response bias, neither our data nor previous studies (Bays et al., 2009; Zhang & Luck, 2008) showed significant bias effects. Consistent with our hypothesis, the CD condition showed significantly lower SD, i.e., higher memory precision, than the ICD condition (Figure 2A, paired sample t-test, t_(32)_ = 4.468, p < 0.001, Cohen’s d = 0.778). While CD trials always exhibited a monotonic increase or decrease in both features (potentially providing mnemonic information), we conducted a critical control analysis to determine whether this property alone could explain the memory advantage. We identified: (1) all ICD trials showing monotonic changes in both domains and (2) their matched CD counterparts sharing identical first two items. Notably, the CD condition retained its performance advantage even in this subset (Figure 2B, paired sample t-test, t_(32)_ = 2.127, p = 0.041, Cohen’s d = 0.370). This indicates that participants specifically leveraged the consistent directional structure—the identical inter-item feature-change ratios across domains—to form compressed memory representations rather than storing each feature dimension independently.

**Figure 2.**
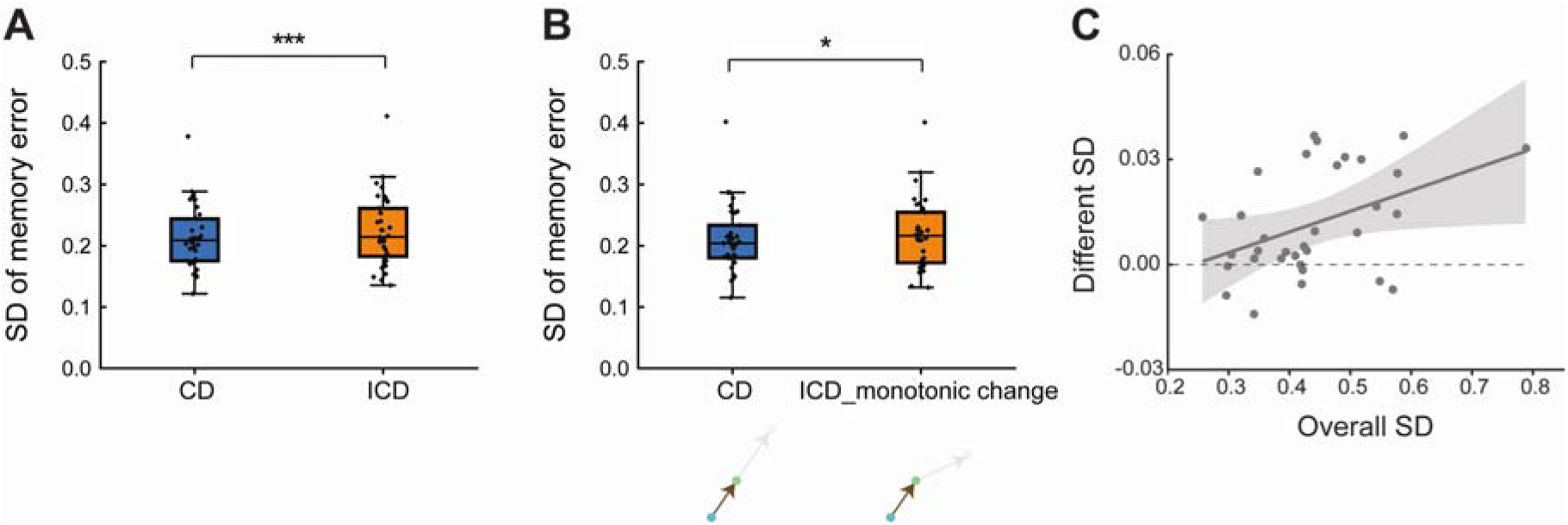
Behavioral performance. (A) Memory performance (standard deviation (SD) of the memory error) for CD (blue) and ICD (orange) conditions (all trials). The horizontal line in the boxplots denotes the median; box outlines denote the 25th and 75th percentiles; whiskers denote 1.5 × the interquartile range. (*: p < 0.05; ***: p < 0.001). (B) Same as A, but for the subsets of trials in the CD condition and ICD condition in which the sequential stimuli showed continuous increase or decrease in both domains. (C) Scatterplot of overall SD (average across the two conditions) (X-axis) and different SD (difference between the two conditions)(Y-axis).

Working memory capacity is considered a significant predictor of general intelligence (Conway et al., 2003), suggesting that individuals may employ distinct strategies to hold a large amount of information in WM. Here, we examined whether participants with high memory capacity (indexed by smaller memory error) were more or less likely to exploit the consistent directional structure for information compression, as reflected in the memory difference between the two conditions. To test this idea, we operationalized WM capacity using overall SD, with a lower SD value indicating higher capacity. As shown in Figure 2C, higher-capacity participants showed a smaller difference between the two conditions (r = -0.437, p = 0.011), which suggested that they may have employed similar mental organization to store multiple items across both conditions. One possible explanation for this finding could be that for high-capacity participants, the task demands may not have approached their WM limits, allowing them to maintain all items without relying on the directional structure for compression. However, lower-capacity participants, struggling to retain all information, may have depended more heavily on the directional structure to compress information within their constrained capacity.

### Neural representation of individual items and directional structure during encoding

The behavioral results suggested that the consistent directional structure in the CD condition led to a behavioral advantage. To investigate its neural implementation, we first focused on the event-related fields (ERFs) across conditions. Given that trials from the two conditions were pseudorandomly interleaved (with conditions distinguishable only at the 3^rd^ item presentation), ERF amplitudes were significantly stronger for the CD condition specifically during the 3^rd^ item presentation (see supplementary figure 3 for the all-sensor and topographic cluster results). We next applied time-resolved representational similarity analysis (RSA) (Kriegeskorte et al., 2008) to examine neural representations at multiple levels, ranging from item-specific information to abstract directional structure. Specifically, each item (1^st^, 2^nd^, and 3^rd^) was characterized as a single point in 2D feature space, allowing the construction of item-specific representational dissimilarity matrix (RDM) via pairwise Euclidean distances based on the 2D feature space between all trials. For the directional structure, we constructed a separate RDM where trial pairs were binary-coded (0: same direction, 1: different direction). Therefore, we had four model RDMs in total (three item-specific and one structural). The neural RDM was generated similarly by calculating pairwise Euclidean distances between multivariate neural response patterns. Through partial correlation analysis, we assessed each model RDM’s unique relationship to neural RDM while controlling for shared variance with other models (Figure 3A, see Methods).

**Figure 3.**
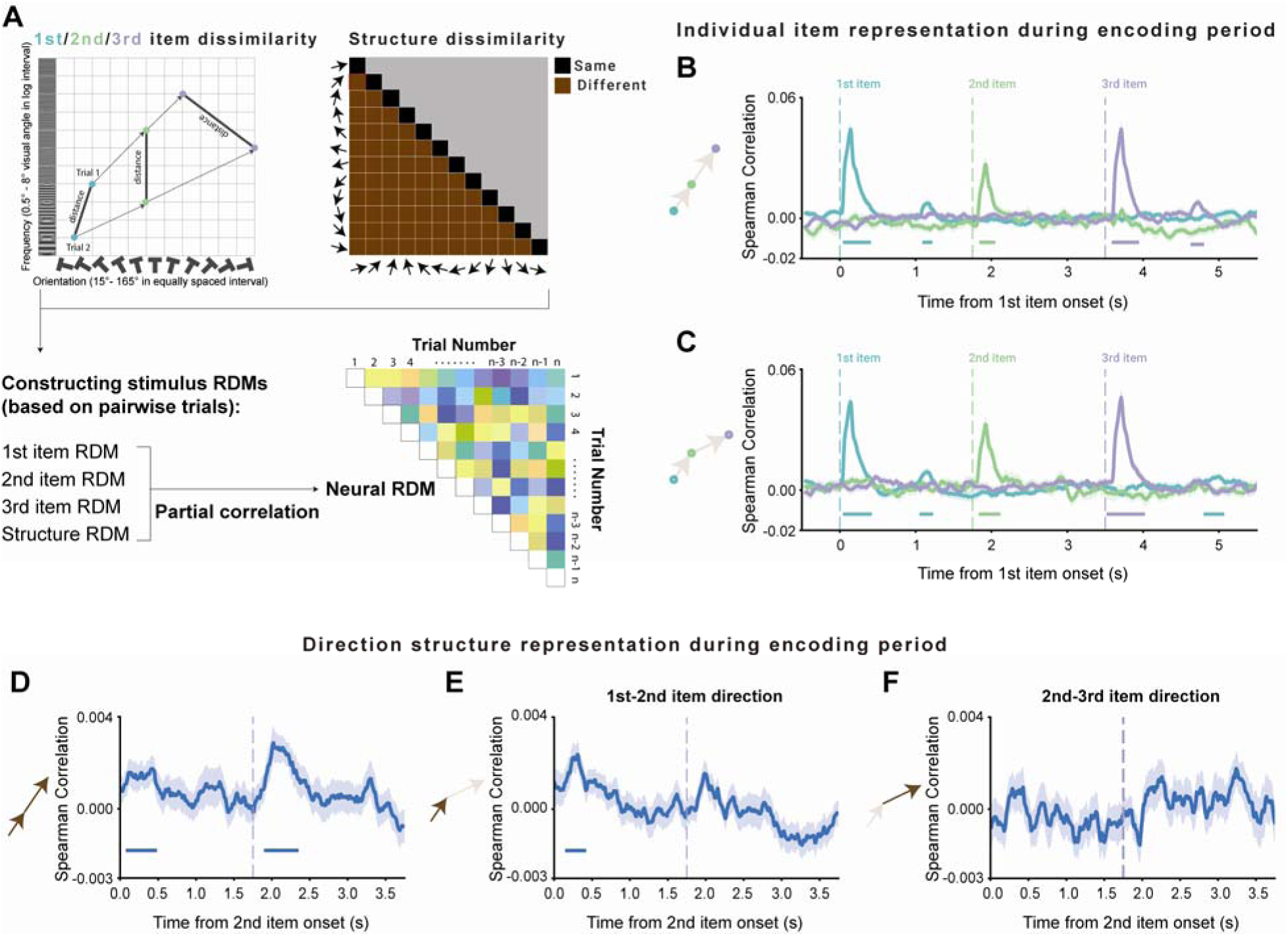
Neural decoding during memory encoding. **(A)** Analysis logic for evaluating the contribution of different aspects of WM contents to neural activity. Individual item RDM was generated by calculating pairwise Euclidean distances based on 2D feature space between all trials. Meanwhile, the directional structure RDM was formed by assessing whether the directions of pairwise trials were the same or different. Neural RDM was estimated by calculating Euclidean distances of the neural responses between pairwise trials. Partial correlation was then applied to measure the association of each stimulus RDM to the neural RDM. **(B)** Time courses of grand average (mean ± SEM) decoding performance for 1^st^ (blue), 2^nd^ (green), and 3^rd^ (purple) items during the encoding period in the CD condition. **(C)** Same as B, but for the ICD condition. **(D)** Time course of grand average (mean ± SEM) decoding performance for the direction information during the encoding period in the CD condition. **(E)** Same as in D, but for the direction from the 1^st^ to 2^nd^ item in ICD condition. **(F)** Same as E, but for the direction from the 2^nd^ to 3^rd^ item in ICD condition. (Horizontal solid line: cluster-based permutation test, cluster-defining threshold p < 0.05, corrected significance level p < 0.05.)

As shown in Figure 3BC, neural representations of all three items were robustly encoded in both conditions. In the CD condition, significant representation clusters emerged following each item’s presentation (1^st^ item: 0.04 – 0.41 s, 1.09 – 1.22 s; 2^nd^ item: 1.84 – 2.05 s; 3^rd^ item: 3.61– 3.92 s, 4.65–4.78 s; cluster p < 0.05). The ICD condition exhibited similar temporal patterns (1^st^ item: 0.04 – 0.42 s, 1.05 – 1.23 s, 4.80 – 5.07 s; 2^nd^ item: 1.83 – 2.12 s; 3^rd^ item: 3.54 – 4.00 s; cluster p < 0.05). Both conditions showed significant encoding of the 1^st^-to-2^nd^ item directional structure during the 2^nd^ item presentation (CD: 0.1–0.46 s post-onset; ICD: 0.17–0.40 s post-onset; cluster p < 0.05). However, consistent with ERF results, the two conditions diverged markedly during the 3^rd^ item presentation period. Specifically, only the CD condition showed significant directional representation (note: the 1^st^-to-2^nd^ item and 2^nd^-to-3^rd^ item directions are identical in this condition; 0.17–0.58 s post-onset, cluster p < 0.05; Figure 3D). In contrast, the ICD condition showed no significant representation of either the 1^st^-to-2^nd^ item (Figure 3E; cluster p > 0.1) or 2^nd^-to-3^rd^ item direction (Figure 3F; cluster p > 0.1). Taken together, these results reveal condition-specific directional encoding profiles.

### Dissociated neural representation patterns during maintenance

We next applied the same RSA analysis to investigate how these multiple information components were held during the maintenance period. Our analysis specifically targeted the period following PING stimulation, a well-established paradigm for probing latent WM representations (Huang et al., 2021; Wolff et al., 2017). Interestingly, the CD condition displayed significant reactivation for directional structure (Figure 4A, 0.47–0.59 s, cluster p < 0.05), but not individual items (Figure 4B, only the 1^st^ item showed marginal reactivation at a relatively late stage, 0.87–0.95 s, 0.05 < cluster p < 0.1). Conversely, the ICD condition revealed the opposite pattern: significant item reactivation (Figure 4D: 1^st^ item, 0.63–0.79 s, cluster p < 0.05; marginal 0.14–0.23 s, 0.05 < cluster p < 0.1) but no directional structure representation (Figure 4C). Note that although the 2^nd^ item reactivation following the 1^st^ item was not significant overall (cluster p > 0.1), high-performing participants showed significant 2^nd^ item reactivation (cluster p < 0.05, see below).

**Figure 4.**
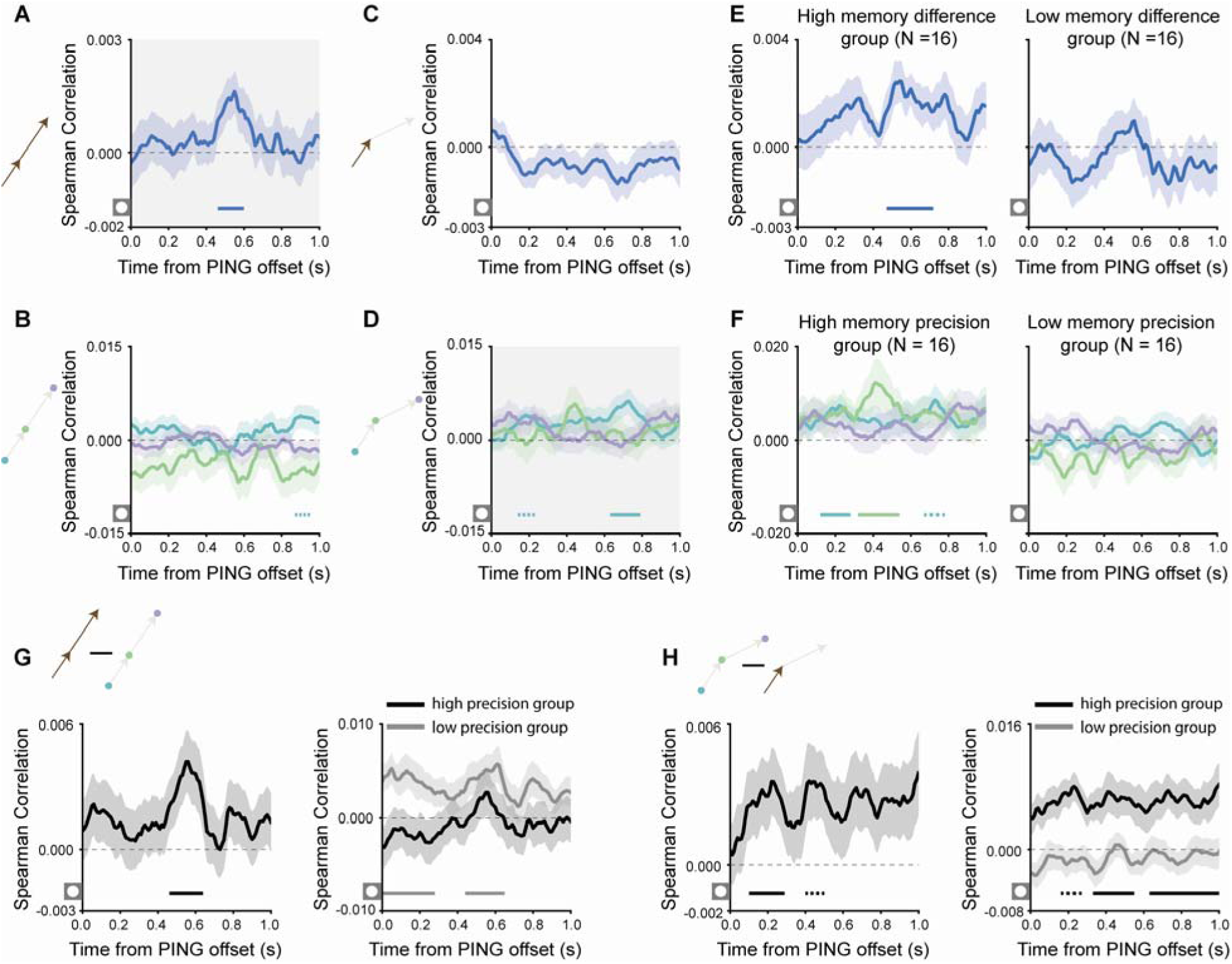
Neural decoding during memory maintenance. **(A)** Grand average (mean ± SEM) decoding performance time course for the direction from the 1^st^ to 2^nd^ items after the PING in the CD condition. **(B)** Grand average (mean ± SEM) decoding performance time courses for 1^st^ (blue), 2^nd^ (green), and 3^rd^ (purple) items after the PING in the CD condition. **(C)** Same as A, but for the ICD condition. **(D)** Same as B, but for the ICD condition. **(E)** Participants were divided into two groups based on memory performance difference between the two conditions. Left panel: grand average (mean ± SEM) direction decoding performance time course in the CD condition for the high memory difference group. Right panel: low memory difference group. **(F)** Participants were divided into two groups based on memory performance in the ICD condition. Left panel: Grand average (mean ± SEM) direction decoding performance time course in the ICD condition for the high memory performance group. Right panel: low memory performance group. **(G)** Left panel: grand average (mean ± SEM) decoding difference between direction and item information (average across three items) in the CD condition. Right panel: participants were divided into two groups based on memory performance in the CD condition. Dark and grey lines separately indicated decoding differences for high and low-performance groups. **(H)** Left panel: same as the left panel of G, but showing the opposite subtraction in the ICD condition. Right panel: participants were divided into two groups based on memory performance in the ICD condition. Dark and grey lines separately indicated decoding differences for high and low-performance groups. (Horizontal solid line: cluster-based permutation test, cluster-defining threshold p < 0.05, corrected significance level p < 0.05. Horizontal dashed line: cluster-based permutation test, cluster-defining threshold p < 0.05, corrected significance level 0.05 < p < 0.1.)

To localize these dissociations, we complemented this all-sensor analysis with searchlight mapping (see Methods). In the CD condition, directional representation was most pronounced over frontal-temporal sensors (0.21–0.65 s, cluster p < 0.05; Figure 5A), with no significant cluster observed for item-specific representation. Conversely, the ICD condition exhibited a widely distributed sensor pattern supporting the 1^st^ item representation (0.01–0.86 s, cluster p < 0.05), peaking in the occipital-parietal sensors (Figure 5B). We focused on the 1^st^ item as it showed the strongest representation (see supplementary figure 5A for similar patterns with averaged three-item representations). No significant cluster emerged for directional structure representation in the ICD condition. Time courses of decoding performance for the identified clusters were shown in supplementary figure 5BC.

**Figure 5.**
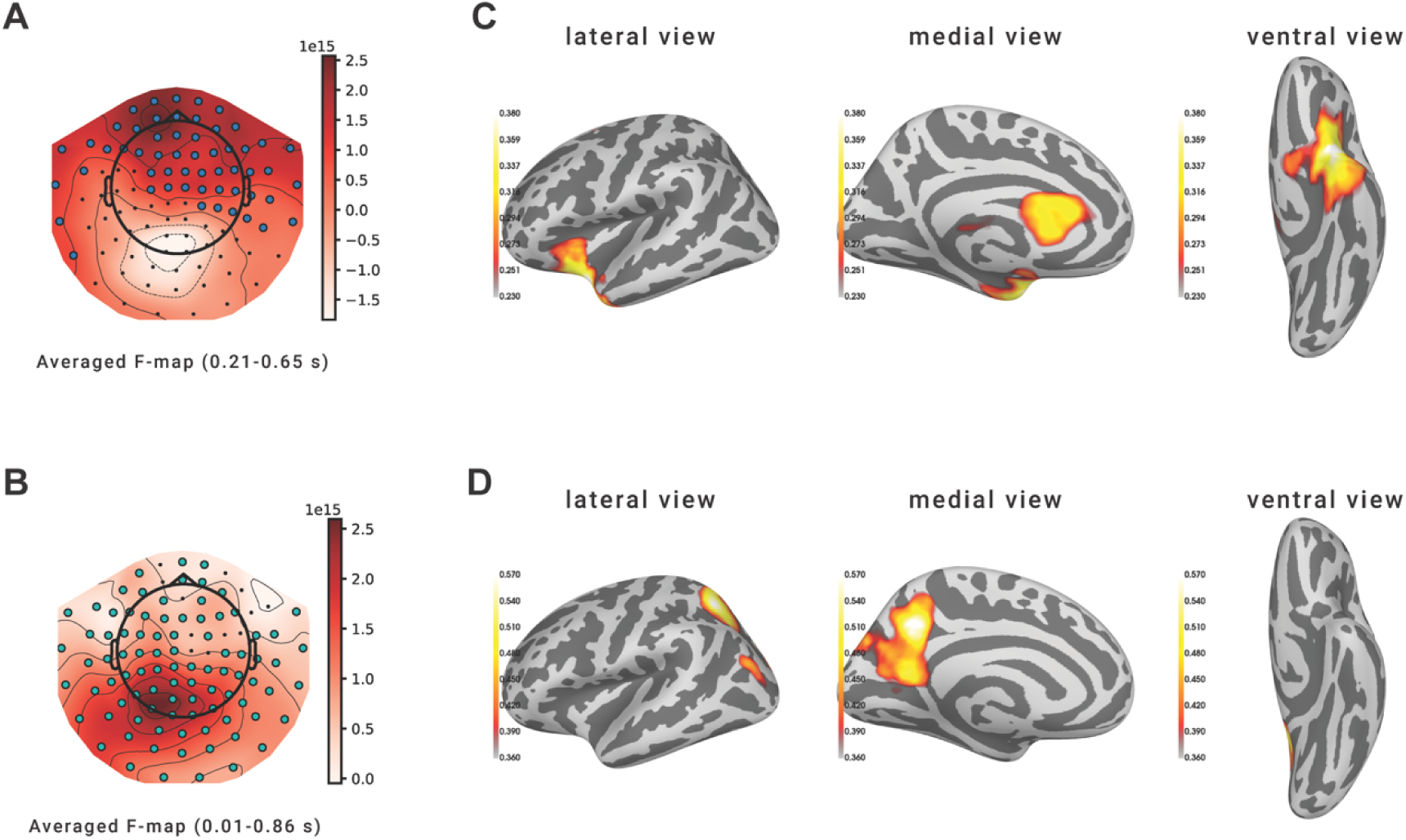
Origin of the dissociated representation patterns. **(A)** Topography plot of significant clusters of direction representation at the sensor level. **(B)** Same as A, but for the 1^st^ item representation. **(C)** Significant brain reactivation for direction information based on lateral, medial, and ventral view. **(D)** Same as C, but for the 1^st^ item representation.

Taken together, these dissociated neural patterns support the efficient coding theory, which posits that neural systems optimize representations according to environmental statistics under capacity constraints (Attneave, 1954). Specifically, when the three items were not independent and could be summarized by directional regularity (CD condition), the brain preferentially represented the abstract directional structure – the most efficient encoding given the stimulus statistics. Conversely, for the statistically independent items (ICD condition), participants relied on the individual item representations rather than directional structures, as in the latter, information compression was impossible and would have demanded additional computational effort.

### Behavioral relevance of the dissociated neural representation patterns during maintenance

Moreover, we examined whether these dissociated patterns had behavioral relevance. As noted previously, the enhanced memory performance in the CD compared to ICD condition reflected the degree to which consistent directional structure was utilized for efficient multi-item compression. To quantify this relationship, we performed a median split (N = 16 per group) based on the memory performance difference between conditions. Notably, only the high memory enhancement group exhibited significant directional structure reactivation (0.47–0.71 s, cluster p < 0.05), while the low-enhancement group showed no significant reactivation (no cluster) (Figure 4E). This finding confirmed that leveraging consistent directional information facilitated efficient multi-item storage. For the ICD condition, to explore behavioral links to the sequential reactivation pattern, we divided participants based on ICD-specific memory performance (Figure 4F). The high-performance group displayed a clear sequential reactivation pattern (0.12–0.28 s (1^st^ item), 0.32–0.54 (2^nd^ item), cluster p < 0.05; 0.67–0.78 s (1^st^ item), 0.05 < cluster p < 0.1), while low-performance group showed no significant reactivation (no cluster). Consistent with our previous work (Huang et al., 2018; Huang & Luo, 2024; Huang et al., 2021), these results underscore the functional role of spontaneous sequential reactivation in memory consolidation.

To investigate how different levels of WM information, specifically, sensory details vs. abstract structure, were optimally represented, we directly compared the two types of representations within each condition. The CD condition exhibited stronger directional structure representation (Figure 4G: left panel, 0.47–0.63 s, cluster p < 0.05), whereas the ICD condition showed dominance of sensory item representation (Figure 4H: left panel, 0.11–0.28 s, cluster p < 0.05; 0.40–0.50 s, 0.05 < cluster p < 0.1). Subsequently, to assess the behavioral relevance of these prioritized representation patterns in different contexts, we grouped participants separately via median splits based on condition-specific memory performance. Importantly, in the CD condition, only the low-performance group relied more heavily on structure representation than on item-specific representation (averaged across items) (Figure 4G: right panel, 0–0.27 s, 0.45– 0.64 s, cluster p < 0.05). This finding aligns with previous behavioral correlations, indicating that participants with low memory capacity were more likely to use consistent directional structure to compress multiple items to fit WM capacity. Meanwhile, in the ICD condition, predominant item representation occurred only in the high memory performance group (Figure 4H: right panel, 0.34–0.54 s, 0.64–1.0 s, cluster p < 0.05; 0.16–0.27 s, 0.05 < cluster p < 0.1). This implies that prioritizing individual item representation is more effective when stimuli lack compressible structure.

### Source origins of the dissociated neural representation patterns during maintenance

Finally, we asked where in the brain the dissociated patterns originated. To identify the cortical sources exhibiting directional structure and item representations, we performed a source reconstruction technique (minimum-norm estimates, MNE) to localize MEG activity during maintenance period. Then we repeated the RSA analysis in the source space (see Methods).

Consistent with the sensor-level results, in the CD condition, we identified a significant directional structure representation in the left anterior temporal and medial frontal cortices with the peak localized around the temporal pole, but found no cluster for individual item representation. Conversely, in the ICD condition, the 1^st^ item representation was localized to the left precuneus and parietal cortex with the peak activation in the superior parietal cortex, while no clusters were observed for directional structure representation. Time courses of decoding performance across source points of clusters were shown in supplementary figure 5DE. Taken together, these findings demonstrate that dissociated brain areas are preferentially engaged in various aspects of WM information—sensory details in unstructured contexts (ICD) versus abstract relational patterns in structured contexts (CD)—aligned with the principles of efficient neural coding.

## Discussion

A fundamental constraint of human cognition is the limited neural resources available for processing and remembering information. However, real-world information is typically statistically structured and predictable. Efficient coding theory proposes that the brain optimizes its neural representations by adapting to these environmental statistics. To test this framework, we designed a sequential multi-item memory task that manipulated statistical regularities among stimuli defined within a 2D feature space. Enhanced memory performance was observed in the consistent direction condition (CD) compared to the inconsistent direction condition (ICD), which indicates that consistent direction is spontaneously extracted to compress multiple information. Interestingly, this benefit was most pronounced in individuals with lower WM capacity. Neural analyses revealed robust reactivation of direction structure during memory maintenance in the CD condition, which correlated with behavioral improvements. However, in the ICD condition where items lacked compressible regularity, individual item information was predominantly represented, with high-performing participants exhibiting clearer sequential activation patterns. These dissociated neural signatures originated from distinct brain regions: directional structure representation localized to the anterior temporal and medial frontal cortex, while item-specific representation emerged in the posterior parietal and precuneus regions.

To overcome the limited capacity in WM, behavioral research over the past decade has consistently demonstrated that the brain exploits environmental statistical regularities to form compressed, efficient memory representations (Brady et al., 2009, 2011; Kaiser et al., 2015; Lew & Vul, 2015; Mathy & Feldman, 2012). However, we know little about the neural mechanisms enabling such regularity-driven compression in WM. In our previous study, by manipulating the consistency of trajectory distance between location and color sequences, we showed that the shared common trajectory structure is strongly encoded and results in a spontaneous neural replay of color sequence during the location recalling period (Huang & Luo, 2024). Building on these findings, the present study investigates how different abstraction levels of information (sensory stimulus vs. abstract structure) are optimally prioritized in dissociated brain areas during memory maintenance period. We propose that this prioritization depends on the availability of compressible statistical regularities, offering novel insight into the neural implementation of optimal coding principles in WM.

Meanwhile, spontaneous reliance on consistent direction information emerged exclusively in participants with low memory capacity. This finding suggests that efficient compression mechanisms are preferentially engaged when memory systems are challenged by high informational demands. According to resource rationality theory, agents optimize behavior by balancing computational costs (e.g., mental effort) against expected rewards, given finite neural resources (Griffiths et al., 2015; Lieder & Griffiths, 2020). For participants with low capacity, the cost of compressing information by extracting statistical regularities (e.g., collapsing 2D feature space into a single domain via consistent directional structure) is outweighed by the prohibitive cost of maintaining all items independently. Conversely, in the ICD condition, where no compressible regularity exists, maintaining items independently incurs lower computational costs than attempting to extract a non-existent regular structure. This capacity-dependent trade-off perfectly illustrates the neural implementation of optimal coding principles, where cognitive systems dynamically balance representational fidelity against computational demands based on both environmental structure and intrinsic capacity limitations.

Notably, unlike typical associative learning or episodic memory paradigms that explicitly train relational structures in either physical or conceptual space (Bao et al., 2019; Constantinescu et al., 2016; Menghi et al., 2025; Simone et al., 2021; Theves et al., 2019), our participants spontaneously extracted latent directional regularities without instruction. This automatic optimization reflects a fundamental biological principle: neural systems naturally evolve to maximize task performance while minimizing a biologically relevant cost (Ganguli & Simoncelli, 2014; Griffiths et al., 2015; Olshausen & Field, 1996). Given that the natural environment exhibits rich statistical structures, efficient memory coding should leverage these regularities to compress information. Our finding—that such statistical knowledge is stored in the temporal and frontal cortex for compression during WM maintenance—further supports the view that WM and long-term episodic memory are interdependent systems (Atkinson, 1968; Cowan, 1988; Daume, Kamiński, Salimpour, et al., 2024; Nichols et al., 2006), dynamically integrated to enable adaptive behavior.

During WM maintenance, stimulus-specific representations are distributed across a cortical network, encompassing sensory, parietal, and frontal cortices (Christophel et al., 2018; Christophel et al., 2017; Ester et al., 2015; Freedman et al., 2001; Harrison & Tong, 2009; Serences et al., 2009; Sprague & Serences, 2013; Yu & Shim, 2017; Zhang & Yu, 2024). Our study extends prior work by demonstrating not only the involvement of the sensory parietal cortex in item-specific representation but also identifying a critical role of the anterior temporal cortex in maintaining abstract directional structures in WM. This result aligns with the recent evidence implicating that medial temporal lobe in WM under conditions of high cognitive demand, such as when managing distractors (Axmacher et al., 2010; Jeneson & Squire, 2012), integrating spatial context (Libby et al., 2014), or maintaining conceptual information (Kamiński et al., 2017). Recent intracranial recordings further reveal that the medial temporal lobe may act as a neural hub bridging WM and long-term memory systems (Daume, Kamiński, Salimpour, et al., 2024). Note that although making definitive anatomical conclusions with MEG is challenging, the observed dissociated representation patterns for abstract directional information and sensory stimulus help reduce the likelihood of false positive findings. Furthermore, while the medial temporal lobe involvement cannot be directly localized with the surface model, its contribution can be inferred indirectly through the cortical surface reactivation patterns, consistent with recent intracranial evidence (Daume, Kamiński, Salimpour, et al., 2024; Daume, Kamiński, Schjetnan, et al., 2024).

The medial temporal lobe, particularly the hippocampal-entorhinal system plays a central role in constructing cognitive maps—structured neural representations that organize knowledge to support adaptive behavior (Hafting et al., 2005; O’Keefe & Dostrovsky, 1971; Tolman, 1948). Originally conceptualized as frameworks for spatial navigation (e.g., enabling route planning or novel shortcuts), cognitive maps have since been formalized as mechanisms for organizing abstract knowledge across domains, facilitating generalization and rapid inference in non-spatial contexts (Behrens et al., 2018; Bellmund et al., 2018; Epstein et al., 2017; O’Keefe, 1978; Tolman, 1948; Whittington et al., 2020). Our findings advance this framework by suggesting that cognitive maps may also operate as efficient coding systems, spontaneously extracting statistical regularities within conceptual spaces to compress information and overcome capacity limitations. This compression mechanism highlights how the brain optimizes WM maintenance by leveraging abstract structure in knowledge representation.

In conclusion, our findings reveal that the brain optimizes WM representations by prioritizing information based on environmental statistics. This process is mediated by dissociated neural systems (anterior temporal for abstract structure, parietal for item details), with low-WM-capacity individuals relying more on structural compression to offset resource limitations. This adaptive mechanism aligns with the principles of efficient coding. Future work could exploit this compression hierarchy framework (structure vs. items) to advance brain-inspired computing or clinical tools. Additionally, exploring how medial temporal lobe-mediated WM and long-term memory integration supports real-world learning and decision-making under resource constraints may bridge gaps in understanding cognitive adaptability.

## Materials and methods

### Participants

Thirty-six participants (aged 18 to 35 years) were recruited via the internal database recruitment system of Max Planck Institute for Human Cognitive and Brain Sciences in Leipzig. The study was approved by the local Ethics Committee of Leipzig University, Germany (Ethics Approval: Nr 018/23-ek). The participants gave written informed consent before the experiment and were compensated for their participation with 12 Euros an hour for the MEG scanning sessions. All participants had normal or corrected-to-normal vision, with no history of psychiatric or neurological disorders. The sample size was determined by a behavioral pilot study using a similar experimental design, which revealed an effect of Cohen’s d = 0.58 with a sample size of 33 participants. Two participants were excluded because they didn’t complete both sessions. Additionally, one participant was excluded from the analysis due to excessive head movement during the scan.

### Stimuli

Sequential grating stimuli were displayed on a rear-projection screen (placed at a viewing distance of 90 cm) with a spatial resolution of 1280 X 1024 pixels and a refresh rate of 60 Hz. Given the circular nature of the orientation, we placed a bar (1.4° X 6° visual angle) above the grating (6° X 6° visual angle), ensuring it remained orthogonal to the stripes of the grating. This setup allowed the grating to represent orientations ranging from 0° to 360°, with a maximum angular difference of 180°. The actual orientation of each grating was drawn from 15° to 165° in a linear interval. The stripe frequency was drawn from 0.5 ° to 8 ° visual angle in logarithmically spaced intervals. In other words, each stimulus can be described as a certain point in a 2D conceptual space defined by frequency and orientation. Participants received instructions about frequency and orientation ranges and became acquainted with both. In the 2D feature space, there were 12 possible directions (ranging from 15° to 345° in 30° increments), representing the ratio of the relative changes (between items) across the two domains.

### Procedure

At the beginning of each trial, three gratings with different frequencies and orientations were sequentially presented for 1 s each at the center of the screen, with 0.75 s intervals between them (Figure 1). After a 1 s delay, a high-luminance white disk was briefly presented to perturb the hidden memory network (Huang et al., 2021; Wolff et al., 2017). Participants then maintained the three gratings in memory for an additional 1 s before receiving instruction about which domain (frequency or orientation) and which specific item they needed to recall. Finally, a test stimulus appeared at the center of the screen. Participants were asked to reproduce only the cued domain (either orientation or frequency) of the target item, by using the mouse wheel to increase or decrease the value of the test item to match the target item as precisely as possible, without any time limits. Note that the non-cued domain of the test stimulus remained identical to that of the target item.

To investigate how multiple items of information are organized based on the underlying regularities, we manipulated abstract direction consistency in the 2D conceptual space. Specifically, in the consistent direction condition (CD), the three sequential items were arranged in a straight line, meaning that the direction of vector from the 1^st^ to 2^nd^ item was identical to that from the 2^nd^ to 3^rd^ item. This allowed the 2D feature space to be compressed into a 1D relationship after extracting consistent directional information, thereby reducing storage demands. In contrast, in the inconsistent condition (ICD), the three items were completely independent, with different directions between 1^st^-to-2^nd^ and 2^nd^-to-3^rd^ items. Therefore, multiple information can’t be effectively compressed as it requires remembering two independent directions. To eliminate the confounding factor of stimulus distribution, the two conditions had the same 1^st^ and 2^nd^ items. This matching ensured that for each ICD trial, there was a corresponding CD trial with the same 1^st^ and 2^nd^ items. This design enabled a direct comparison of how identical stimuli were differently organized and stored in WM under different contexts. Besides, the starting point (1^st^ item) and the inter-item distance were varied, with the constraint of a minimum angular distance of 15° and frequency distance of 0.2773° visual angle in logarithmic scale to make items distinguishable. This ensured that the three items were fully sampled from the 2D continuous feature space (see supplementary figure 1). Crucially, the two conditions were interleaved such that they could only be distinguished following the presentation of the 3^rd^ item. Before the MEG sessions, participants were asked to complete 120 practice trials to familiarize themselves with the task. During MEG recordings, they needed to complete 720 trials in total, which were divided into 2 sessions. Each session took around 2.5 hours (including breaks).

### MEG recordings

Participants completed the two MEG sessions inside a sound-attenuated and magnetically shielded room. Neuromagnetic data were acquired using a 306-sensor MEG system (204 planar gradiometers, 102 magnetometers, MEGIN Vectorview system) at the Max Planck Institute for Human Cognitive and Brain Sciences, Leipzig, Germany. For the preparations, we first placed 5 head-position indicator (HPI) coils on the participant’s head: 3 were on the forehead while the other 2 were on the left and right mastoid bone behind the ears. We then used an electromagnetic digitizer system to digitize the head position by 3 fiducial anatomical markers (the nasion, left, and right preauricular points) and the 5 HPI coils. Finally, we digitized at least 600 points on the scalp to obtain the whole shape of the head. By utilizing head digitalization, the MEG head position can be spatially aligned with individual structural MRI images for source reconstruction analysis.

### MEG preprocessing

The MEG data were recorded at 1000 Hz sampling frequency and applied an online low-pass filter (cutoff frequency 330 Hz). The MEG data were preprocessed offline following the FLUX pipeline (Ferrante et al., 2022) (https://www.neuosc.com/flux) using MNE-Python (version 1.3.1). Specifically, the spatiotemporal signal space separation (tSSS) was used to remove external noise (Taulu & Simola, 2006). Muscle activity was detected and removed. Independent component analysis was then performed in each subject to remove eye-blink and heartbeat artifact components, and the remaining components were then back-projected to channel space. After preprocessing, the data were epoched into segments, band-pass filtered between 2 and 30 Hz, and downsampled to 100 Hz. To further identify artifacts, the variance (collapsed over channels and time) was first calculated for each trial. Trials with excessive variances were removed. MEG data were baseline-corrected before further analysis. Specifically, the time range from 200 ms to 100 ms before the presentation of the 1^st^ item in each trial was used as a baseline to be subtracted.

### Data analysis

#### Behavioral performance analysis

For each orientation, the response error was quantified as the difference between the reported orientation and the true target orientation in each trial. A similar analysis was applied to the frequency domain, except that both the reported and actual frequencies were converted to a logarithmic scale. Memory performance was estimated by calculating the standard deviation of response error separately for frequency and orientation, which reflects the reciprocal of memory precision. Thus, a lower value indicates better memory performance. We then combined the two standard deviations by first normalizing them with corresponding ranges and then summing them (see separate results for each domain in supplementary figure 2). As mentioned earlier, since the two conditions held the same 1^st^ and 2^nd^ items, we combined their memory and focused on the difference between the two conditions (see separate results for each item in supplementary figure 2).

#### Source modeling

Structural T1-weighted images were acquired using a 3T Siemens Prisma scanner with either a 32-channel coil or a 64-channel coil. One participant was unable to complete the anatomical scan, so the FreeSurfer standard template (fsaverage) (Dale et al., 1999) was used for the forward model. MEG source modeling was conducted using dynamic statistical parametric mapping (dSPM) (Dale et al., 2000), which is based on depth-weighted minimum-norm estimates (MNE) (Hämäläinen & Ilmoniemi, 1994). This analysis was applied to epoched data that had been baseline-corrected from −0.2 s to −0.1 s relative to stimulus onset. To construct the forward model, the MRI images were manually aligned with the digitized head shape. For the participant without MRI scan, the Freesurfer template was warped to match their digitized head shape and thus creating a subject-specific template. Using MNE-Python, a single shell Boundary Elements Model (BEM) was constructed based on the inner skull surface obtained from FreeSurfer. A surface-based forward model covering the whole brain was then generated. The lead field matrix was calculated based on the head’s position relative to the MEG sensor array. The noise covariance matrix for the baseline period (−0.5 s to 0 s before the stimulus onset) was computed based on all sensors and combined with the forward model to generate a common spatial filter. Finally, this spatial filter was applied to the baseline-corrected epoch data to estimate the source of the MEG signals.

#### Time-resolved RSA analysis

To investigate how different aspects of information, ranging from sensory details to abstract direction structure, were held in memory, we applied Representational Similarity Analysis (RSA) (Kriegeskorte et al., 2008) to the baseline-corrected sensor data and source estimates. The analysis was performed using the MNE-Python RSA Toolbox (mne-rsa toolbox, version 0.10, https://users.aalto.fi/~vanvlm1/mne-rsa/). For each item, we created representational dissimilarity matrices (RDM) by computing pairwise Euclidean distances between all trial pairs. Before Euclidean distance calculations, frequency values were converted to a logarithmic scale, and then both frequency and orientation values were normalized by dividing them by their corresponding ranges. The RDM for direction structure was constructed by evaluating whether trial pairs shared identical or different directional structures.

Neural RDMs were derived from spatiotemporal neural patches. At the sensor level, patches were constructed for each sensor by aggregating neighboring sensors within a 5cm spatial radius (centered on that sensor) and a 50 ms temporal window. In source space, they were defined by source estimates within a spatial radius of 2 cm and a temporal radius of 50 ms. For each patch, pairwise Euclidean distances between neural patterns across trials were computed to generate neural RDMs. Next, for each patch, the contribution of all these model RDMs (3 item RDMs + 1 direction RDM) to neural RDM was estimated by running partial correlations. This measured the association between each model RDM and the neural RDM while controlling for the influence from the remaining model RDMs (Figure 3A). The results in Figures 3&4 reflect averages across all brain-wide patches. Qualitatively similar (but weaker) patterns were observed when using a neural RDM derived from all gradiometer sensors (see supplementary figure 4). To localize neural representation (i.e., searchlight mapping), we additionally performed spatiotemporal cluster-based permutation tests (10000 permutations) on the estimated correlation maps. Statistical significance was thresholded at p < 0.05 (cluster-defining threshold: p < 0.05), to control for multiple comparisons (Figure 5).

#### Statistical analysis

We examined the temporal dynamics of neural decoding performance using cluster-based permutation approach (10000 permutations) (Maris & Oostenveld, 2007). Specifically, for both stimulus presentation and delay periods, we first identified temporal clusters of contiguous significant time points (p < 0.05, two tails for the encoding period, one tail for the maintaining period) using one-sample t-test against chance level (0). We then estimated cluster-level statistics by computing the size of each cluster. Finally, we determined cluster significance probabilities through a Monte Carlo randomization (10000 permutations) procedure to establish a null distribution. For spatiotemporal cluster-based permutation tests in sensor and source spaces, we applied the same permutation testing framework while incorporating spatial adjacency in cluster definition.

## Author contributions

Q.H. and C.F.D. conceived the initial idea. Q.H. designed the experiment and developed the tasks. Q.H. acquired the data. Q.H. planned and performed analyses. Q.H. and C.F.D. discussed the results. Q.H. wrote the initial draft. Q.H. and C.F.D. finalized the manuscript.

## Acknowledgments

This work was supported by Humboldt Research Fellowship for Postdocs to Q.H. and Max Planck Society. We thank Nicholas Menghi, Yangwen Xu, Simone Vigano, Casper Kerren, Alejandro Tabas, Marit Petzka and Muzhi Wang for their helpful comments.

## Supplementary

**Supplementary Figure 1.**
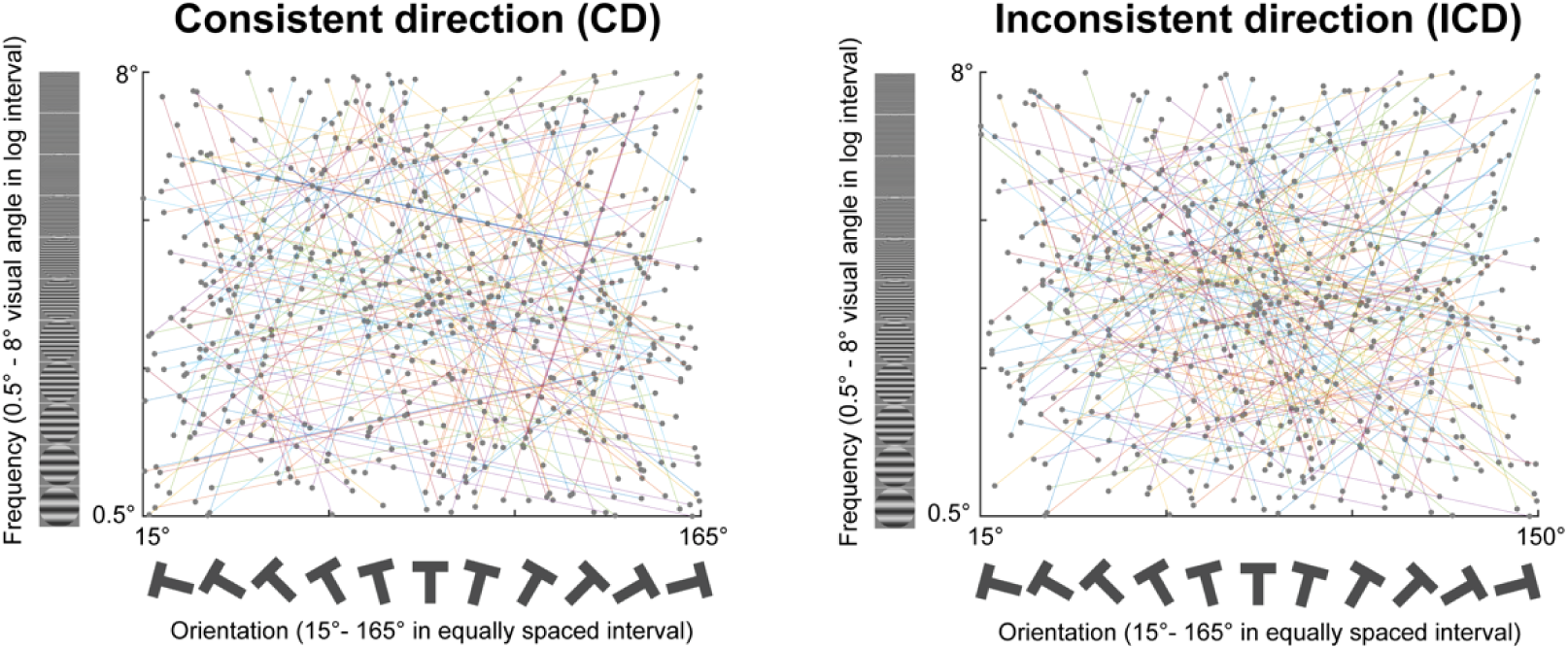
Illustration of how the three sequential items were randomly sampled within the continuous 2D feature space for individual participants in CD (left panel) and ICD (right panel) conditions. One dot indicates one item. Three connected dots (with two lines) denote one trial.

**Supplementary figure 2.**
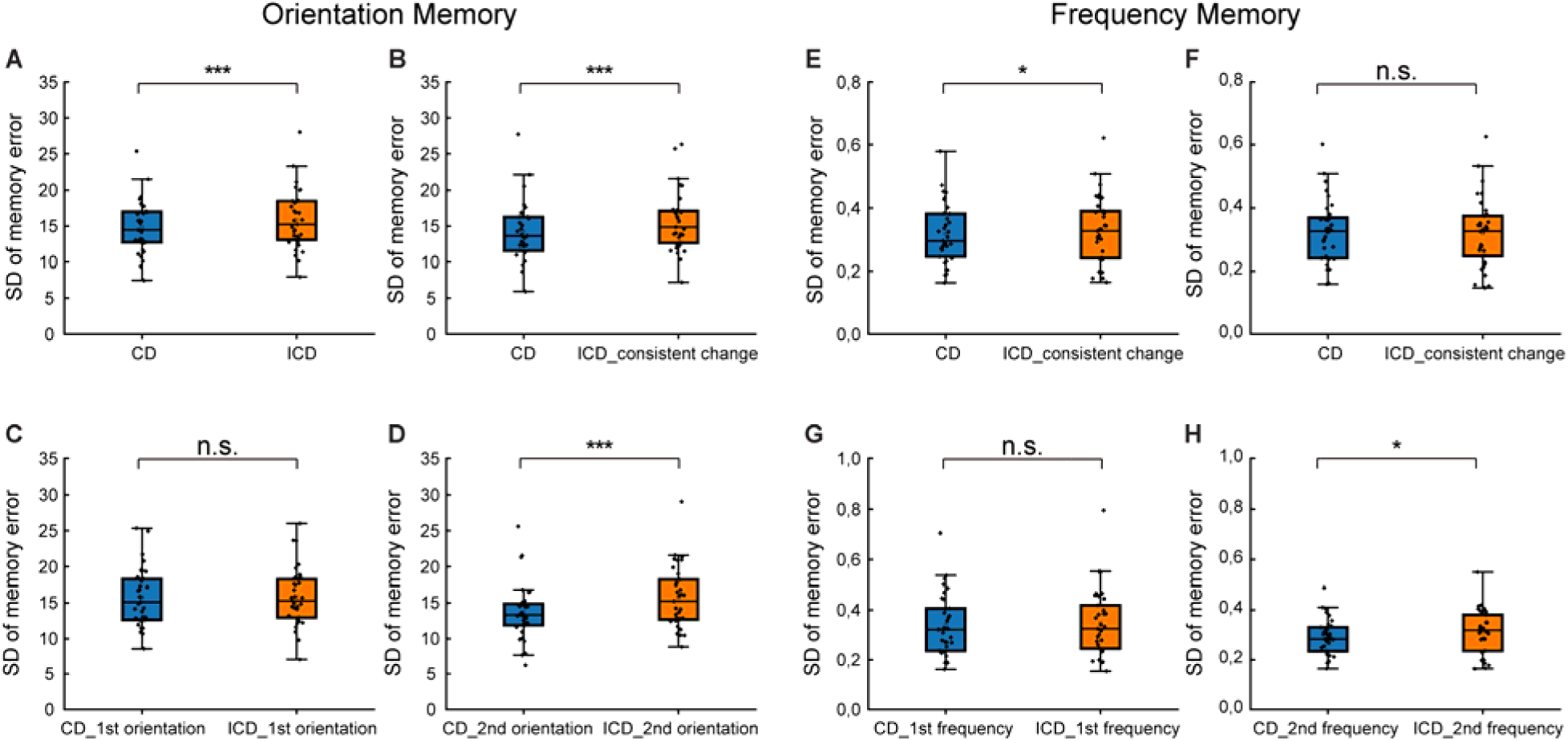
(ABCD) Orientation memory performance. A: memory performance (combined the 1^st^ and 2^nd^ items) shown seperately for CD and ICD conditions. B: Same as A, but selected trials with consistent change in ICD condition, found the paired trials in CD condition and compared them. C: Same as A, but only focused on the 1^st^ item performance. D: Same as A, but only focused on the 2^nd^ item performance. (EFGH) Same analysis configuration as (ABCD), but for frequency memory performance. (*: p < 0.05; ***: p < 0.001, n.s.: p > 0.05).

**Supplementary figure 3.**
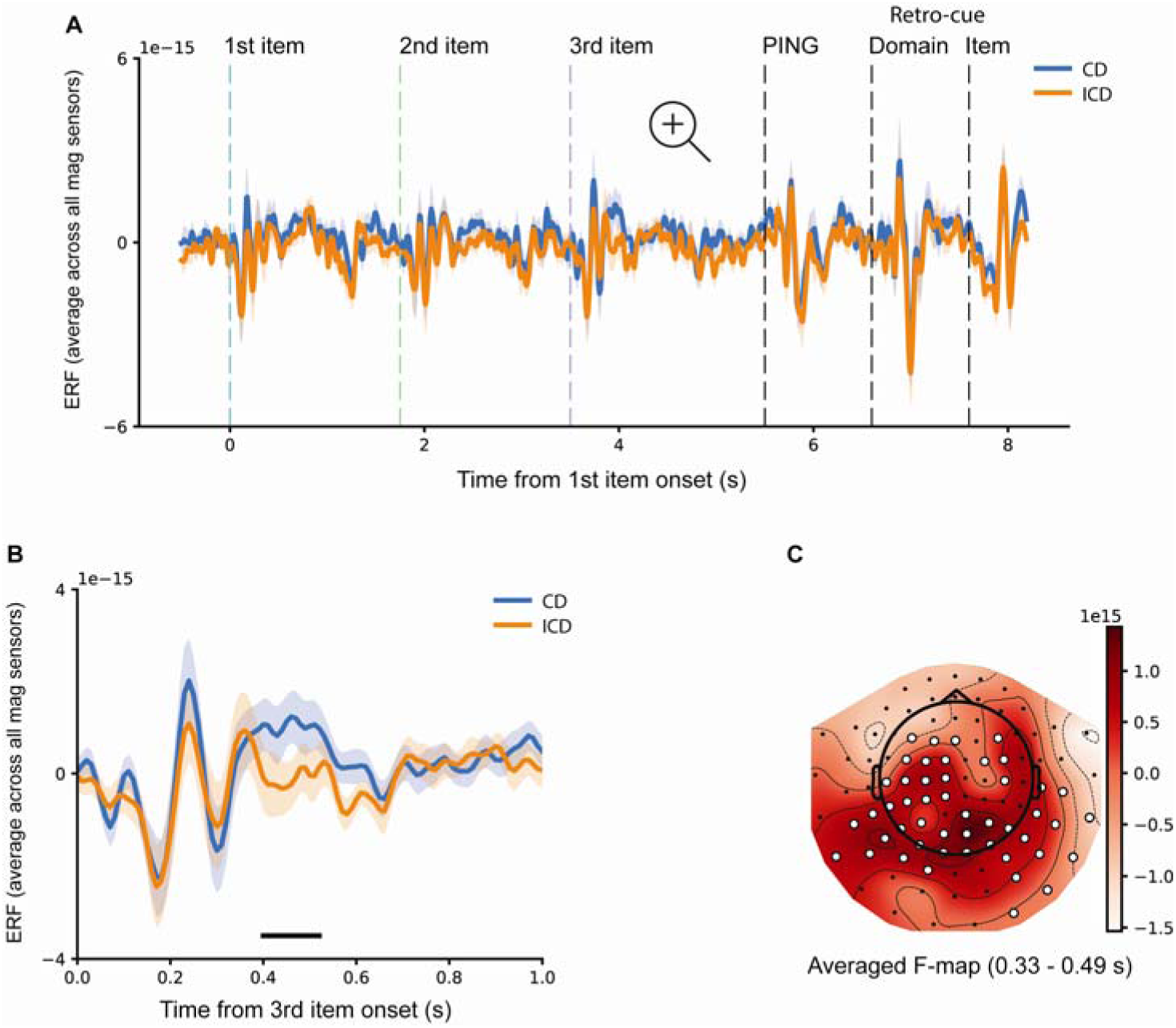
(A) Time courses of grand average (mean ± SEM) event-related potential (ERF) (average across all magnetometers) for CD (blue) and ICD (orange) conditions. (B) Magnified view of the time window corresponding to the 3^rd^ item presentation (same data as A). (C) Topographic distribution of significant cluster from CD vs. ICD ERF comparison during the presentation of the 3^rd^ item. (Horizontal solid line: cluster-based permutation test, cluster-defining threshold p < 0.05, corrected significance level p < 0.05.)

**Supplementary figure 4.**
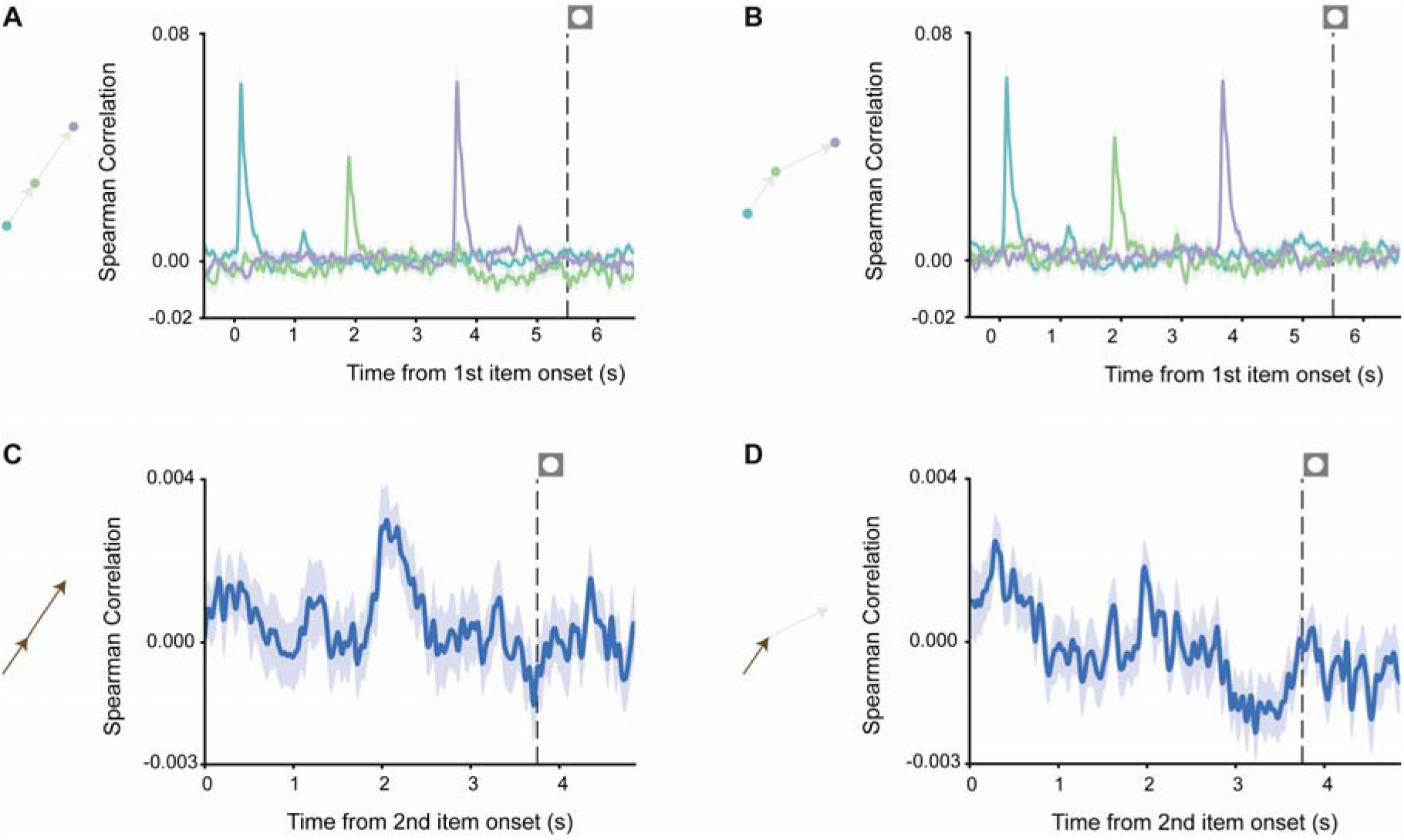
(A) Time courses of grand average (mean ± SEM) decoding performance based on the all gradiometor sensors for the 1^st^ (blue), 2^nd^ (green), and 3^rd^ (purple) items during encoding and maintaining period in the CD condition. The dashed vertical line indicates the onset of the PING stimulus. (B) Same as A, but for the ICD condition. (C) Time course of grand average (mean ± SEM) for directional structure representation after the onset of the 2^nd^ item in the CD condition. (D) Same as C, but for the direction from the 1^st^ to 2^nd^ item in the ICD condition.

**Supplementary figure 5.**
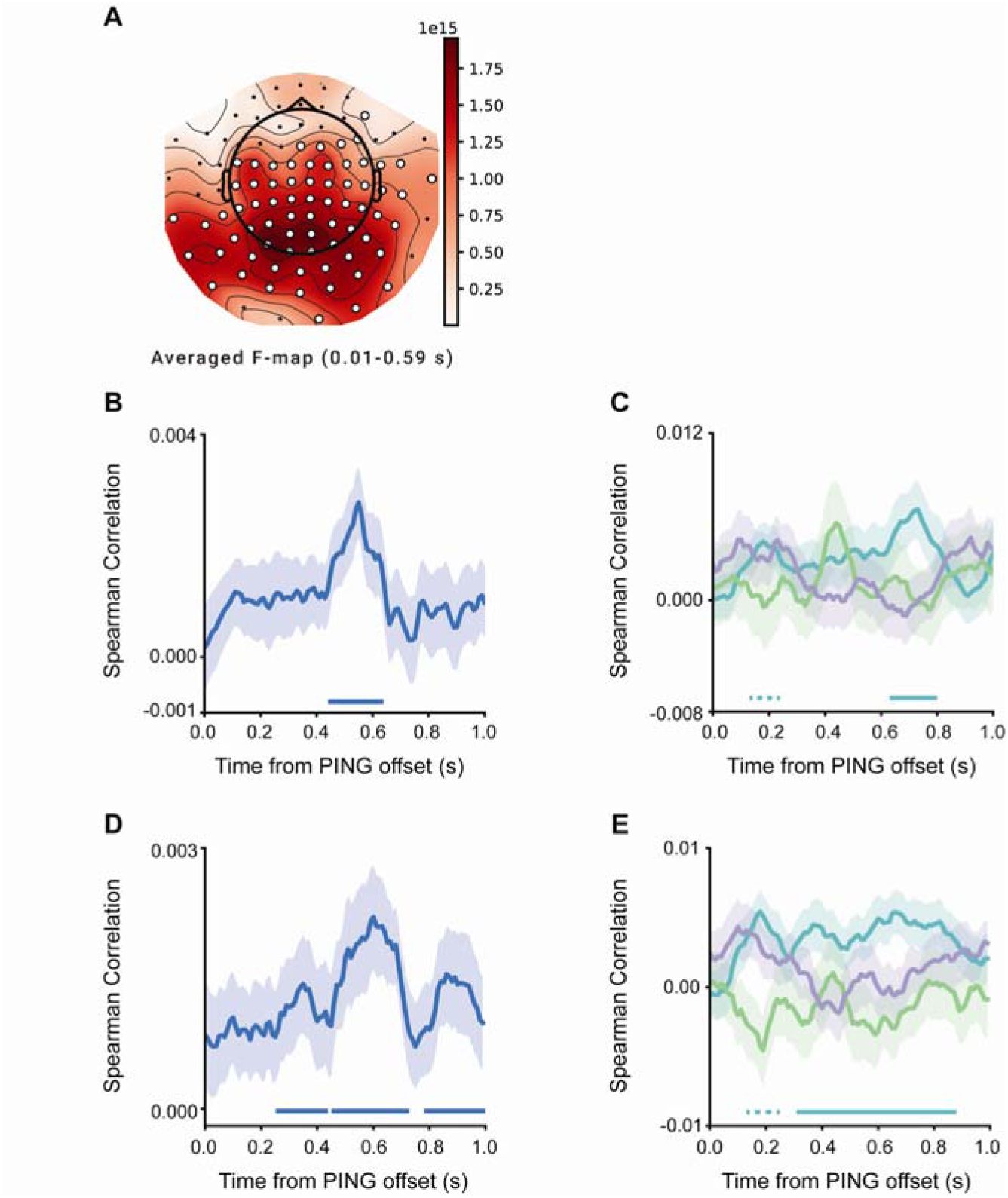
(A) Topographic distribution of significant clusters for item-specific decoding performance (across all three items) during memory maintaining period in the ICD condition. (cluster-defining threshold: p < 0.05, corrected significance: 0.05 < p < 0.1.) (B) Time course of grand average (mean ± SEM) decoding performance for directional information in CD condition across the sensor points of the cluster observed in Figure 5A. (C) Time courses of grand average (mean ± SEM) decoding performance for individual item information in ICD condition across the sensor points of the cluster observed in Figure 5B. (D) Time course of grand average (mean ± SEM) decoding performance for directional information in CD condition across the source points of the cluster observed in Figure 5C. (E) Time courses of grand average (mean ± SEM) decoding performance for individual item information in ICD condition across the source points of the cluster observed in Figure 5D. (Horizontal solid line: cluster-based permutation test, cluster-defining threshold: p < 0.05, corrected significance: p < 0.05. Horizontal dashed line: cluster-based permutation test, cluster-defining threshold: p < 0.05, corrected significance: 0.05 < p < 0.1.)

